# Aldehyde accumulation in *Mycobacterium tuberculosis* with defective proteasomal degradation results in copper sensitivity

**DOI:** 10.1101/2023.02.11.528157

**Authors:** Gina Limón, Nora M. Samhadaneh, Alejandro Pironti, K. Heran Darwin

## Abstract

*Mycobacterium tuberculosis* is a major human pathogen and the causative agent of tuberculosis disease. *M. tuberculosis* is able to persist in the face of host-derived antimicrobial molecules nitric oxide and copper. However, *M. tuberculosis* with defective proteasome activity is highly sensitive to nitric oxide and copper, making the proteasome an attractive target for drug development. Previous work linked nitric oxide susceptibility with the accumulation of *para*-hydroxybenzaldehyde in *M. tuberculosis* mutants with defective proteasomal degradation. In this study, we found that *para*-hydroxybenzaldehyde accumulation was also responsible for copper sensitivity in these strains. We showed that exogenous addition of *para*-hydroxybenzaldehyde to wild-type *M. tuberculosis* cultures sensitized bacteria to copper to a degree similar to that of a proteasomal degradation mutant. We determined that *para*-hydroxybenzaldehyde reduced the production and function of critical copper resistance proteins of the regulated in copper repressor (RicR) regulon. Further, we extended these Cu-sensitizing effects to an aldehyde that *M. tuberculosis* may face within the macrophage. Collectively, this study is the first to mechanistically propose how aldehydes can render *M. tuberculosis* susceptible to an existing host defense and could support a broader role for aldehydes in controlling *M. tuberculosis* infections.

**IMPORTANCE:** *M. tuberculosis* is a leading cause of death by a single infectious agent, causing 1.5 million deaths annually. An effective vaccine for *M. tuberculosis* infections is currently lacking, and prior infection does not typically provide robust immunity to subsequent infections. Nonetheless, immunocompetent humans can control *M. tuberculosis* infections for decades. For these reasons, a clear understanding of how mammalian immunity inhibits mycobacterial growth is warranted. In this study, we show aldehydes can increase *M. tuberculosis* susceptibility to copper, an established antibacterial metal used by immune cells to control *M. tuberculosis* and other microbes. Given that activated macrophages produce increased amounts of aldehydes during infection, we propose host-derived aldehydes may help control bacterial infections, making aldehydes a previously unappreciated antimicrobial defense.

## INTRODUCTION

*Mycobacterium tuberculosis* is a major human-exclusive pathogen and the causative agent of tuberculosis disease (TB). TB is responsible for more than 1 million deaths annually and was the deadliest infectious disease worldwide prior to the SARS-CoV-2 pandemic (https://www.who.int/news-room/fact-sheets/detail/tuberculosis). TB can be cured by lengthy treatments with multiple antibiotics; however, antibiotic-resistant strains of *M. tuberculosis* are increasingly prevalent. Thus, an improved understanding of *M. tuberculosis* pathogenesis and existing host responses to the bacteria are required to aid in the development of new treatment strategies.

Macrophages, a primary cell niche of *M. tuberculosis*, require the production of nitric oxide (NO) for robust resistance to various infections, in particular *M. tuberculosis* (1, 2). NO is a free radical that can form toxic reactive intermediates that damage macromolecules, including nucleic acids, proteins, and lipids [reviewed in (1)]. While the precise mechanism of NO-mediated toxicity to *M. tuberculosis* is unknown, *M. tuberculosis* requires a Pup-proteasome system (PPS) to resist NO (3). In the PPS, numerous proteins destined for degradation are post-translationally modified with Pup (<u>p</u>rokaryotic <u>u</u>biquitin-like <u>p</u>rotein) by the ligase PafA (<u>p</u>roteasome <u>a</u>ccessory <u>f</u>actor <u>A</u>) [reviewed in (4)]. Pup directly binds to a hexameric ATPase, <u>m</u>ycobacterial <u>p</u>roteasome <u>A</u>TPase (Mpa; ARC in non-mycobacteria), which unfolds pupylated proteins and delivers them into a proteasome core protease. In the absence of either Mpa or PafA, numerous proteins fail to be degraded. Importantly, when the PPS is nonfunctional in *M. tuberculosis*, the enzyme lonely guy (Log) accumulates (5). Log plays a role in the production of adenine-based hormones called cytokinins, which are required for normal growth and development in plants (6). *M. tuberculosis* is not found in plants and instead uses cytokinins to induce gene transcription that alters the mycobacterial cell envelope, a phenotype that is not linked to NO resistance (5, 7). In plants, cytokinins are enzymatically broken down into adenine and various aldehydes (8). While it is unknown how cytokinins are metabolized in *M. tuberculosis*, at least one cytokinin-associated aldehyde, *para*-hydroxybenzaldehyde (*p*HBA), measurably accumulates in a PPS mutant, and this aldehyde is sufficient to render wild-type *M. tuberculosis* sensitive to NO (5). However, the mechanistic link between aldehyde accumulation and NO sensitization of *M. tuberculosis* remains unknown.

In addition to NO, there is evidence that macrophages use copper (Cu) to defend against *M. tuberculosis* and other microbial infections (9–12). *M. tuberculosis* has two defined Cu-responsive systems to mitigate Cu toxicity: the CsoR (copper-sensitive operon repressor) operon (13, 14) and the RicR (regulated in copper repressor) regulon (15). CsoR and RicR are paralogues that regulate Cu resistance, although the RicR regulon appears to play a more substantial role in Cu resistance in mice (12, 15–18). When Cu concentrations are low, RicR binds to the promoters of five loci, including its own promoter, blocking their expression. When Cu levels increase, Cu binds to RicR, preventing it from associating with DNA and allowing gene expression. Two RicR-regulated gene products, MmcO, a multicopper oxidase, and MymT, a Cu metallothionein, confer resistance to Cu; deletion of either *mmcO* or *mymT* results in Cu hyper-susceptibility *in vitro* (16, 18, 19). While the deletion of any single RicR-regulated gene does not attenuate *M. tuberculosis* growth in C57BL6/J mice, the production of a “Cu-blind” RicR protein that constitutively represses expression of the entire regulon results in highly Cu-sensitive bacteria that are attenuated for growth *in vivo* (18).

Our lab reported that *M. tuberculosis* PPS mutants are hypersensitive to Cu (18). We reasoned this phenotype may primarily be due to the repression of the RicR regulon in these strains. RicR is not a PPS substrate therefore it is not the accumulation of RicR itself that leads to regulon repression (15). Given that aldehyde accumulation in a PPS mutant sensitizes *M. tuberculosis* to NO, we hypothesized that aldehydes also cause Cu sensitivity. In this work, we determined that a mutation that prevents the accumulation of *p*HBA suppresses the Cu-sensitive phenotype of an *M. tuberculosis* PPS mutant. Additionally, we found *p*HBA can disrupt the production and activities of Cu-binding proteins. Finally, we showed that the aldehyde methylglyoxal (MG), which accumulates in macrophages during *M. tuberculosis* infections, can also reduce *M. tuberculosis* resistance to Cu.

## RESULTS

### Cu sensitivity of an *M. tuberculosis mpa* mutant was suppressed by disruption of *log*

*M. tuberculosis* strains with disruptions in *mpa* or *pafA* (PPS degradation-deficient strains) are hypersensitive to Cu (18) and NO (3). We previously showed that NO hypersensitivity is due to an accumulation of the PPS substrate Log; a transposon insertion in *log* in an *mpa* mutant suppresses its NO-sensitive phenotype (5). We tested if the disruption of *log* could also suppress the Cu-sensitive phenotype of an *mpa* mutant. We confirmed that a *Δmpa::hyg* mutant, hereafter referred to as an *mpa* mutant, is sensitive to Cu (**Fig. 1A**, left panel, center bars; see **Table 1** for all strains used in this work). Disruption of *log* (*log*::MycoMarT7) in the *mpa* mutant strain (hereafter referred to as an *mpa log* mutant) restored wild-type (WT) Cu resistance (**Fig. 1A**, left panel, right-side bars). Given that disruption of *log* prevents *p*HBA accumulation in the *mpa* mutant (5), this result suggested accumulation of *p*HBA in PPS mutants contributes to the Cu-sensitivity of these strains.

**Fig. 1.**
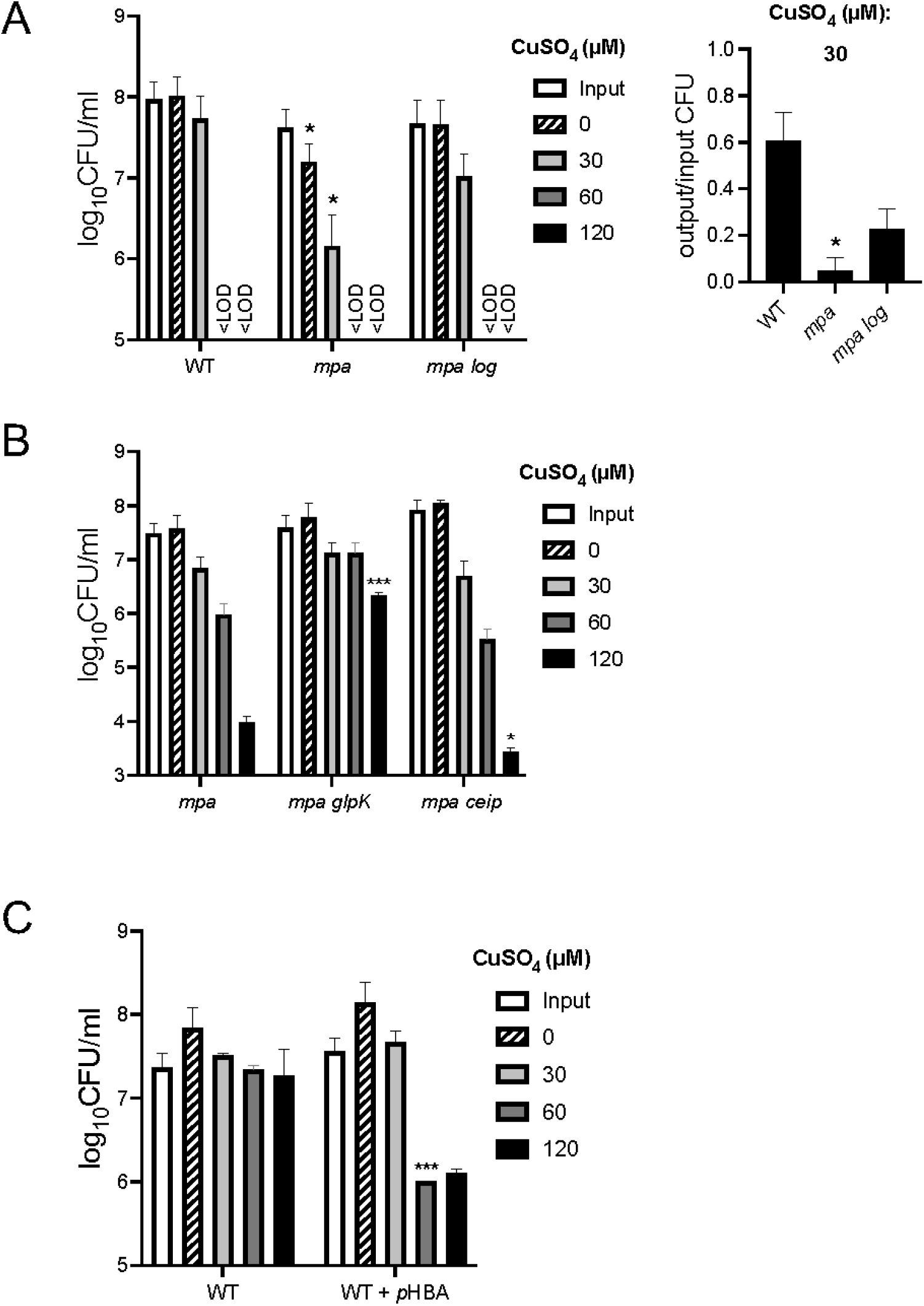
Copper sensitivity of a PPS mutant is suppressed by mutations in *log* or *glpK*. (**A**) Cu sensitivity assay shows a PPS *M. tuberculosis* mutant is hyper-sensitive to Cu. Bacteria were incubated for 10 days in the indicated CuSO_4_ concentrations. “Input” (white bars) indicates CFU at the beginning of the experiment and “0” indicates how many bacteria were present after 10 days of incubation without added CuSO_4_ (striped bars). Bars represent mean with standard deviation (SD). “<L.O.D.” indicates below the limit of detection (100 CFU). Significant differences were calculated comparing CFU of strains to the first strain on the x-axis at the same CuSO4 concentration using an unpaired t-test, with * =*p* < 0.05; *** = *p* < 0.001. Unlabeled bars showed no significant differences. Right panel: Given the slower growth of the mpa mutant, we calculated the ratio of the mean output CFU to mean input CFU for 30 μM CuSO_4_, which more accurately describes the survival of this strain in Cu. (**B**) Cu sensitivity assay performed as in (A) with previously characterized strains suppressed for NO sensitivity. (**C**) Exogenous addition of *p*HBA sensitized WT *M. tuberculosis* to Cu. Cu sensitivity assays performed with WT *M. tuberculosis* incubated with or without 1.2 mM *p*HBA for 24 hours prior to addition of CuSO_4_. For all data shown, experiments are representative of two independent experiments, each done in technical triplicate.

**Table 1.**
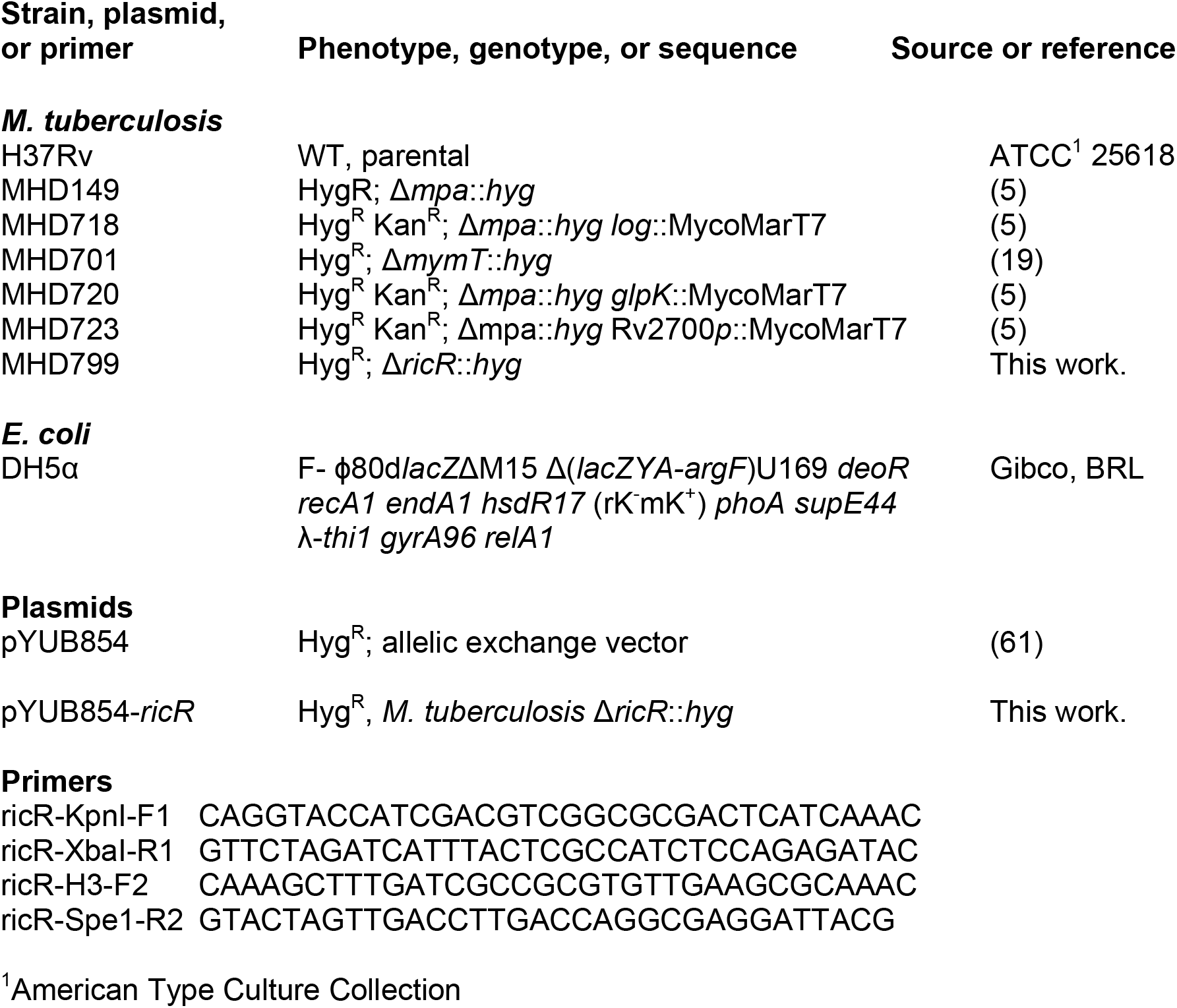
Strains, plasmids, and primers used in this study

Previous work by our lab found two additional mutations that suppress the NO sensitivity of an *mpa* mutant (5). These suppressor mutations are in *glpK*, which encodes a probable glycerol kinase, and in the promoter of *cei*, which encodes a possible secreted alanine-rich protein involved in cell membrane integrity (20). We found the *mpa glpK (Δmpa::hyg glpK::* MycoMarT7) double-mutant had increased Cu resistance compared to the parental *mpa* strain (**Fig. 1B**, center bars). GlpK phosphorylates glycerol, resulting in the production of several aldehydes including MG and glyceraldehyde-3-phosphate. Thus, disruption of *glpK* might reduce the overall aldehyde burden in the *mpa* mutant, mitigating the negative effects of *p*HBA accumulation in this strain. In contrast, the *mpa ceip (Δmpa::hyg ceip::* MycoMarT7) mutant was more sensitive to Cu than the *mpa* mutant (**Fig. 1B**, right side bars). Given that defects in *cei* increase membrane permeability (20) it is possible its reduced expression makes the bacterial cytosol more accessible to Cu.

We next tested if exogenously added *p*HBA could sensitize WT *M. tuberculosis* to Cu. We used 1.2 mM *p*HBA for all experiments in this study because this concentration, which is non-toxic on its own, can robustly synergize with NO to sterilize *M. tuberculosis* cultures (5). We preincubated WT *M. tuberculosis* with *p*HBA in minimal media for 24 hours before exposing the bacteria to Cu, reasoning that pre-treatment with *p*HBA would allow for transcriptional or other changes that affect Cu resistance. Additionally, *p*HBA added concomitantly with CuSO_4_ appeared to inhibit Cu-dependent killing for unclear reasons. As previously reported, *p*HBA alone had no effect on CFUs recovered (**Fig. 1C**, striped bars) (5). Pre-incubation of bacteria with *p*HBA reduced CFU after Cu treatment compared to Cu treatment alone (**Fig. 1C**, compare dark grey and black bars between untreated and *p*HBA-treated). Thus, endogenously-produced or exogenously added *p*HBA sensitized *M. tuberculosis* to Cu.

### Expression of Cu-responsive genes was reduced in *p*HBA-treated *M. tuberculosis*

Microarray analysis of PPS mutants lacking either *mpa* or *pafA* resulted in the identification of five promoters controlled by the DNA binding protein RicR (15). RicR dissociates from DNA in the presence of Cu, leading to the expression of genes required for robust Cu resistance and virulence (15, 18). Another Cu responsive operon, the CsoR operon, is also repressed in PPS mutant strains. Like RicR, CsoR releases repression of its operon after binding to Cu (13, 21). The CsoR operon includes *ctpV*, which encodes a cation transporter that contributes to Cu resistance and virulence (17). Given that PPS mutants are Cu sensitive and have reduced expression of two Cu responsive systems, we hypothesized *p*HBA accumulation in PPS mutants was responsible for the gene expression changes. To test this hypothesis, we performed RNA-Seq on WT *M. tuberculosis* treated with *p*HBA (**Fig. 2** and **Table S1**). We found that of 3,979 genes analyzed, six genes were significantly differentially upregulated more than two-fold, and 35 genes were significantly differentially downregulated by more than two-fold. Remarkably, there was considerable overlap in the gene expression profiles of PPS mutants v. WT *M. tuberculosis* (15) and *p*HBA-treated bacteria vs. untreated WT bacteria (**Table 2** and **Table 3**, gray rows). The substantial overlap in the transcriptomes supported our hypothesis that *p*HBA contributes to the transcriptional signature of PPS mutants. Importantly, all members of the Cu-sensing regulons were down-regulated in PPS mutants and *p*HBA-treated *M. tuberculosis* (**Fig. 2** and **Table S1**).

**Fig. 2.**
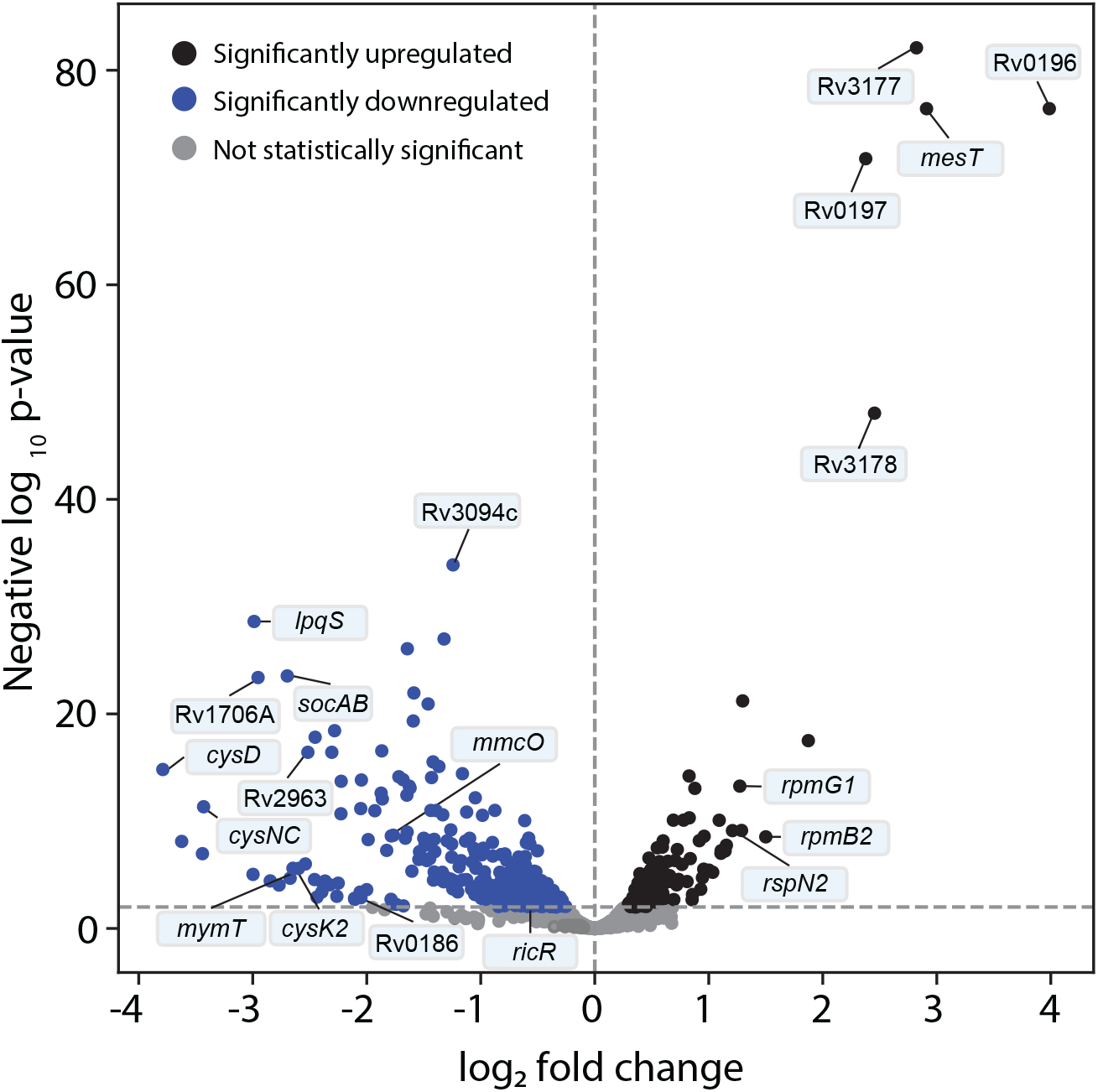
Volcano plot of significantly regulated genes in untreated v. *p*HBA-treated *M. tuberculosis*. x-axis: log_2_-fold change; y-axis: negative log_10_ *p*-value. Dotted horizontal line indicates cutoff for genes to be considered significantly differentially regulated (*p* < 0.01). Dotted vertical line indicates division between up-regulated (black dots) and down-regulated (blue dots) genes. Among the labeled genes are the RicR regulon (Rv0846, *mmcO, lpqS*, Rv1706A, *socAB, mymT, cysK2*, Rv2963, *ricR*) and select members of the Zur regulon (*rpmB2, rpsN2, rpmG1*).

**Table 2.**
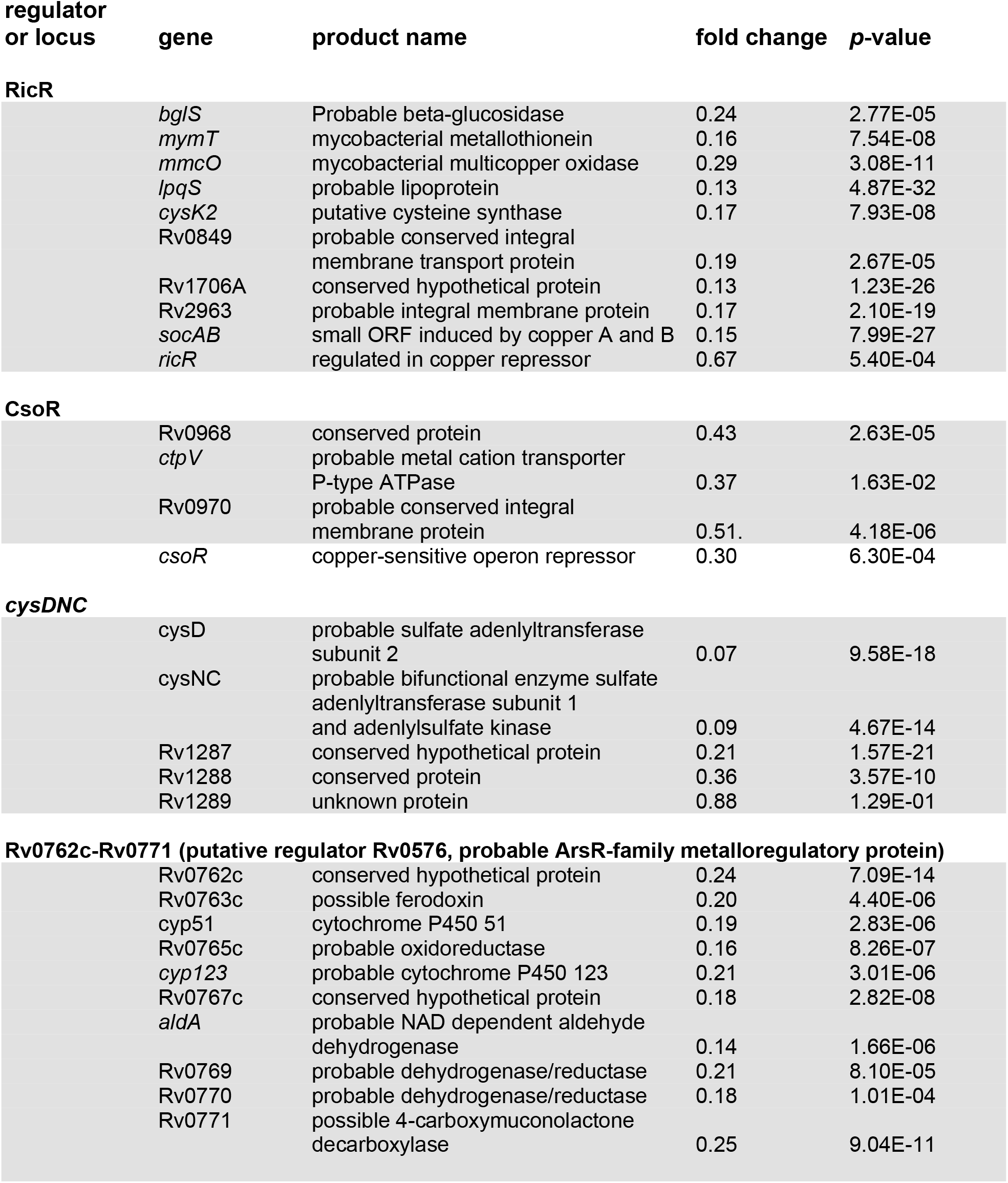
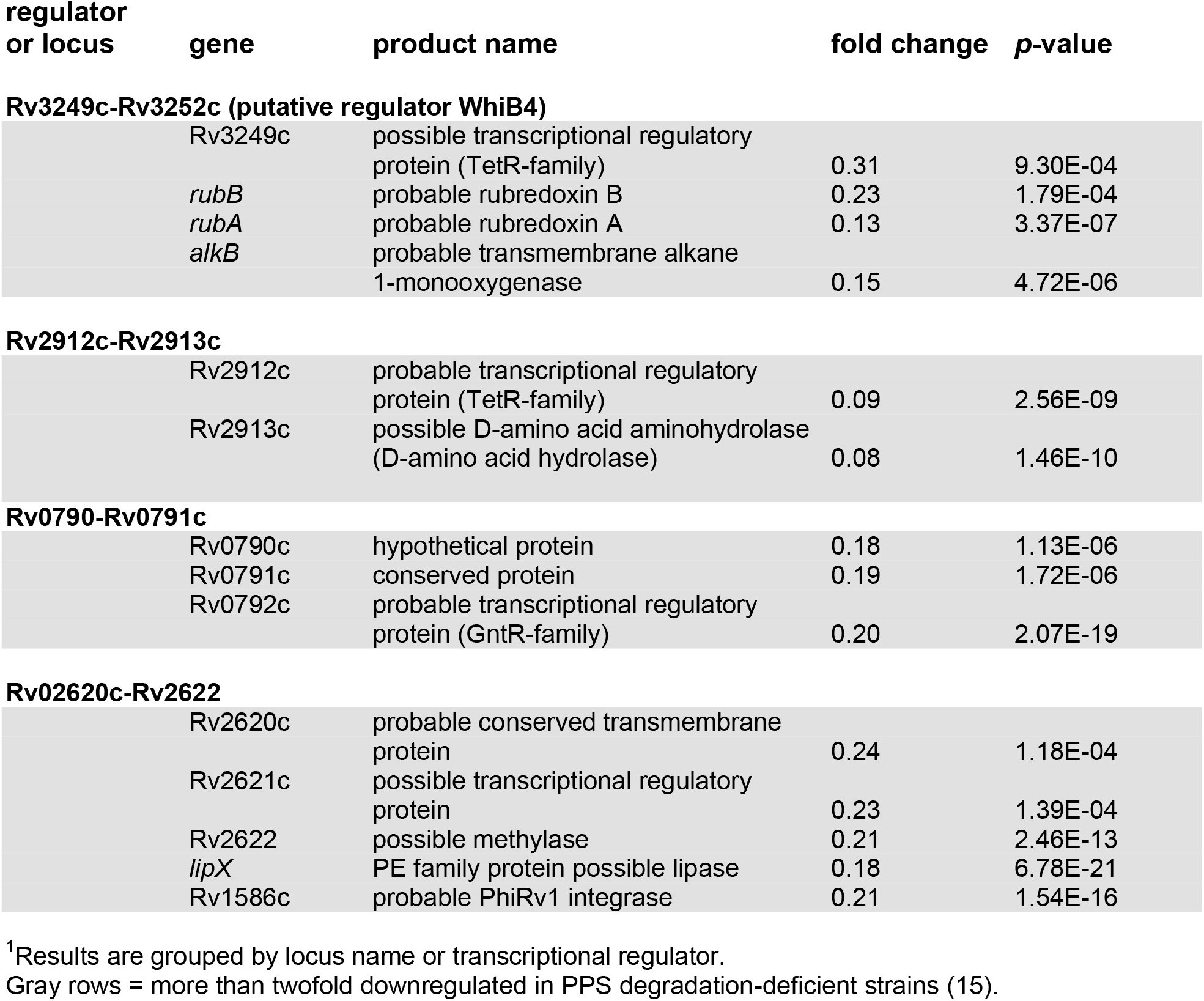
Downregulated operons and regulons in WT *M. tuberculosis* treated with *p*HBA relative to untreated bacteria.

**Table 3.** Upregulated operons and regulons in WT *M. tuberculosis* treated with *p*HBA relative to untreated bacteria.

| regulator<br>or locus | gene | product name | fold change | p-value |
| --- | --- | --- | --- | --- |
| **Rv3173c-Rv3178** |  |  |  |  |
|  | Rv3173c | probable transcriptional regulatory protein (probable TetR/AcR-family) | 3.66 | 1.48E-20 |
|  | Rv3174 | probable short-chain dehydrogenase or reductase | 27.58 | 0 |
|  |  | Rv3175 possible amidase | 2.31 | 1.11E-11 |
|  | *mesT* | probable epoxide hydrolase | 7.52 | 3.85E-80 |
|  | Rv3177 | possible peroxidase | 7.07 | 4.08E-86 |
|  | Rv3178 | conserved hypothetical protein | 5.48 | 1.41E-51 |
| **Rv0195-Rv0197** |  |  |  |  |
|  | Rv0195 | possible two-component transcriptional regulatory protein (LuxR-family) | 2.46 | 2.17E-24 |
|  | Rv0196 | possible transcriptional regulatory protein | 15.86 | 2.99E-80 |
|  | Rv0197 | possible oxidoreductase | 5.19 | 2.21E-75 |
| **Zur** |  |  |  |  |
|  | Rv0106 | conserved hypothetical protein | 2.15 | 2.32E-09 |
|  | Rv0280 | PPE family protein PPE3 | 1.81 | 5.17E-05 |
|  | Rv0281 | Possible S-adenosylmethionine-dependent methyltransferase | 1.53 | 1.38E-04 |
|  | *eccA3* | ESX-3 conserved component type VII secretion system (T7SS) protein | 1.41 | 2.99E-05 |
|  | *eccB3* | ESX-3 T7SS protein | 1.42 | 1.06E-04 |
|  | *eccC3* | ESX-3 T7SS protein | 1.41 | 2.94E-05 |
|  | Rv0286 | PPE family protein PPE4 | 1.19 | 1.04E-01 |
|  | *esxG* | ESAT-6 like protein | 1.27 | 1.69E-02 |
|  | *esxH* | low molecular weight protein antigen 7 | 1.29 | 9.20E-03 |
|  | *espG3* | ESX-3 secretion associated protein | 1.19 | 5.15E-02 |
|  | *eccD3* | ESX-3 T7SS probable transmembrane protein | 1.15 | 1.99E-01 |
|  | *mycP3* | probable membrane-anchored mycosin | 1.21 | 1.89E-02 |
|  | *eccE3* | ESX-3 T7SS secretion system protein | 1.14 | 1.03E-01 |
|  | *rpsR2* | 30S ribosomal protein S18 | 2.00 | 1.18E-07 |
|  | *rpsN2* | 30S ribosomal protein S14 | 2.44 | 1.06E-11 |
|  | *rpmG1* | 50S ribosomal protein L33 | 2.42 | 4.54E-16 |
|  | *rpmb2* | 50S ribosomal protein L28 | 2.83 | 4.58E-11 |
|  | Rv2059 | conserved hypothetical protein | 1.84 | 7.93E-16 |
|  | *zur* | zinc uptake regulator protein | 0.92 | 2.23E-01 |

<sup>1</sup>Results are grouped by locus name or transcriptional regulator.
Gray rows = more than twofold upregulated in PPS degradation-deficient strains (15).

While the RicR regulon encodes eight genes, only two (excluding *ricR*) have been implicated in Cu resistance: *mmcO* and *mymT*, with a *mymT* mutant having the strongest Cu-sensitive phenotype (18, 19). To follow up the transcriptional analysis, we sought to determine if protein levels of either of these key Cu resistance proteins were reduced in *p*HBA treated bacteria or a PPS mutant. Given the low abundance and small size (53 amino acids) of MymT in WT *M. tuberculosis* (15, 19), we instead looked at MmcO. MmcO is a membrane-associated Cu oxidase that is hypothesized to convert Cu(I) into less toxic Cu(II) based on its high similarity to other Cu oxidases (16, 18). We assessed levels of MmcO in Cu-treated *mpa, mpa log*, and *ricR* null (*ΔricR::hyg*) strains relative to a WT parental strain. As previously reported, the *ricR* null mutant over-produces MmcO relative to WT bacteria; in contrast, the *mpa* mutant had approximately two-fold less MmcO relative to WT bacteria (**Fig. 3A**). Consistent with the suppressive effect of the *log* mutation on Cu sensitivity in a PPS mutant (**Fig. 1A**), the *mpa log* strain had WT levels of MmcO (**Fig. 3A**).

**Fig. 3.**
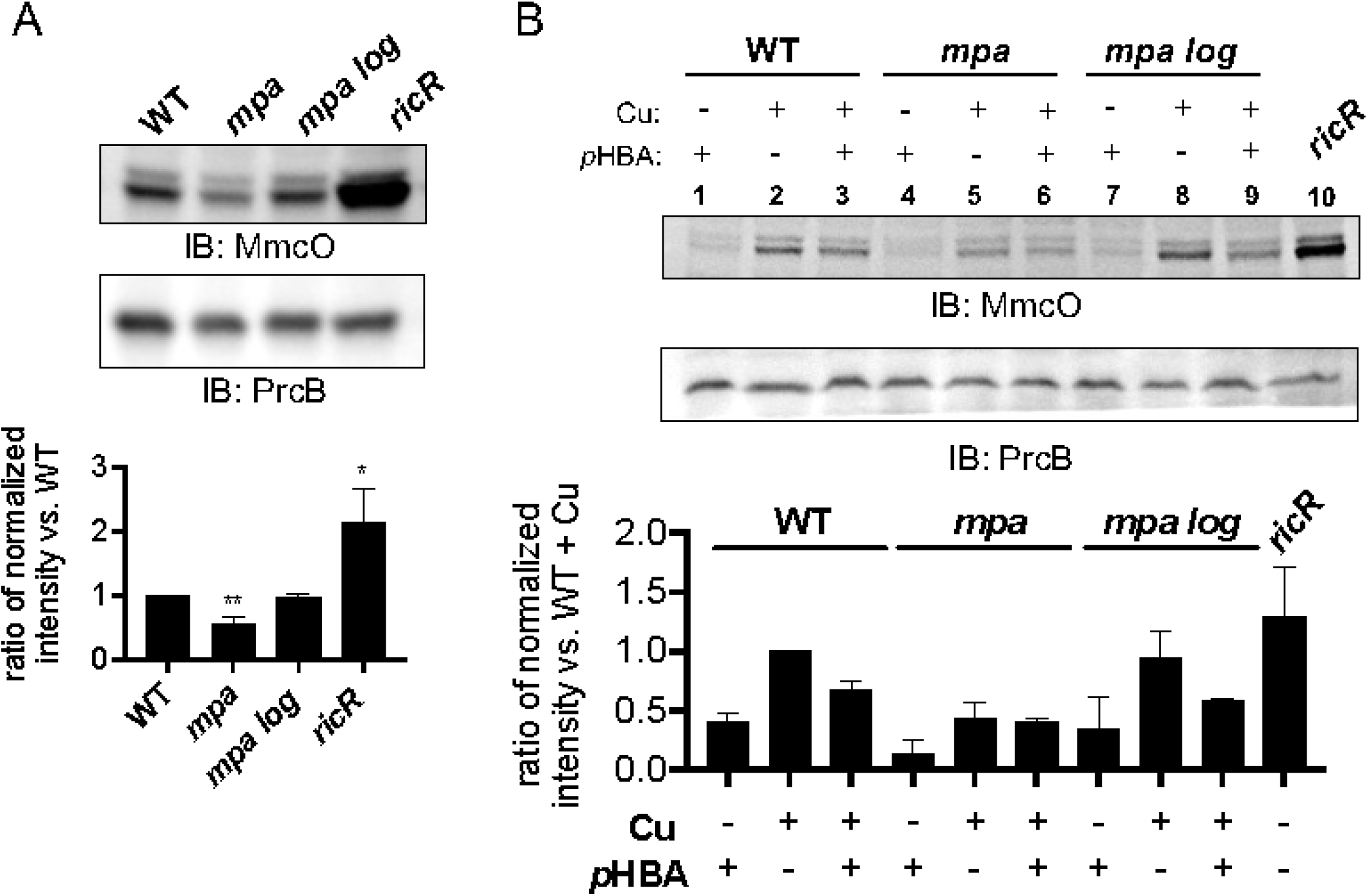
MmcO levels are reduced in a PPS mutant and in *p*HBA-treated *M. tuberculosis*. (**A**) *M. tuberculosis* strains indicated were grown in Sauton minimal media and treated with 50 μM CuSO_4_ for 24 hours prior to collection of equivalent amounts of bacteria for whole cell lysates and preparation for immunoblotting. Proteins were probed using polyclonal rabbit antibodies raised against MmcO or the β-subunit of the proteasome (PrcB) as a loading control. This experiment is a representative of three biological replicates. Below: Quantification of band intensity from three independent experiments with MmcO levels normalized to PrcB levels before calculating the intensity ratio between each strain compared to the WT strain. Bars indicate mean with SD error bars. Significance was calculated using an unpaired t-test to WT with * =*p* < 0.05; ** =*p* < 0.01. Unlabeled bars showed no significant differences. (**B**) Immunoblotting for MmcO in strains treated with or without 1.2 mM *p*HBA for four hours before treatment with CuSO_4_. Below: Quantification as in (A) of MmcO levels normalized to PrcB levels for each strain and condition, relative to MmcO levels in the WT strain treated with Cu only for two independent experiments. Bars indicate mean with SD error bars.

We next determined if exogenously added *p*HBA could affect MmcO levels. In the absence of Cu, *p*HBA treatment alone resulted in low MmcO levels in all strains, comparable to MmcO levels in untreated *M. tuberculosis* (**Fig. 3B**, lanes 1, 4 and 7) (16, 18). Cu treatment of the WT strain increased MmcO levels as previously reported (16, 18) (**Fig. 3B**, lanes 1 v. 2), whereas *p*HBA reduced levels of MmcO produced in Cu-treated WT bacteria by approximately one-third (**Fig. 3B**, lanes 2 v. 3). Even with the addition of Cu, the *mpa* strain showed low levels of MmcO, nearly half the level observed in Cu-treated WT bacteria (**Fig. 3B**, lanes 2 v. 5), and the addition of *p*HBA had a minor effect on the already low MmcO levels in the *mpa* mutant (**Fig. 3B**, lanes 5 v. 6). MmcO levels in the *mpa log* strain were similar to that of the WT strain (**Fig. 3B**, lanes 2 v. 8, and lanes 3 v. 9). Together these results indicate that the presence of endogenously produced or exogenously added *p*HBA reduced the levels of at least one RicR-regulated gene product, which supports our transcriptional data.

### *p*HBA altered the function of the major Cu resistance protein, MymT

Up to this point, our data support a model whereby PPS-defective *M. tuberculosis* are hyper-susceptible to Cu due to a reduction in Cu-responsive gene expression caused by *p*HBA. However, we could not rule out the possibility that *p*HBA directly disrupts the function of one or more of the Cu-responsive proteins. A way to test this hypothesis is to measure the effect *p*HBA has on a *ricR* null mutant, which constitutively expresses high levels of all of the RicR regulon genes. We performed a Cu-sensitivity assay on a *ricR* mutant pretreated with *p*HBA. Because *ricR* mutants are highly resistant to Cu (15), we needed to use a higher Cu concentration than in previous experiments to kill the bacteria. At the highest CuSO_4_ concentration used, *p*HBA robustly sensitized the *ricR* mutant to Cu (**Fig. 4A**, black bars), suggesting that repression of RicR-regulon gene expression was not the only mechanism by which *p*HBA sensitized *M. tuberculosis* to Cu.

**Fig. 4.**
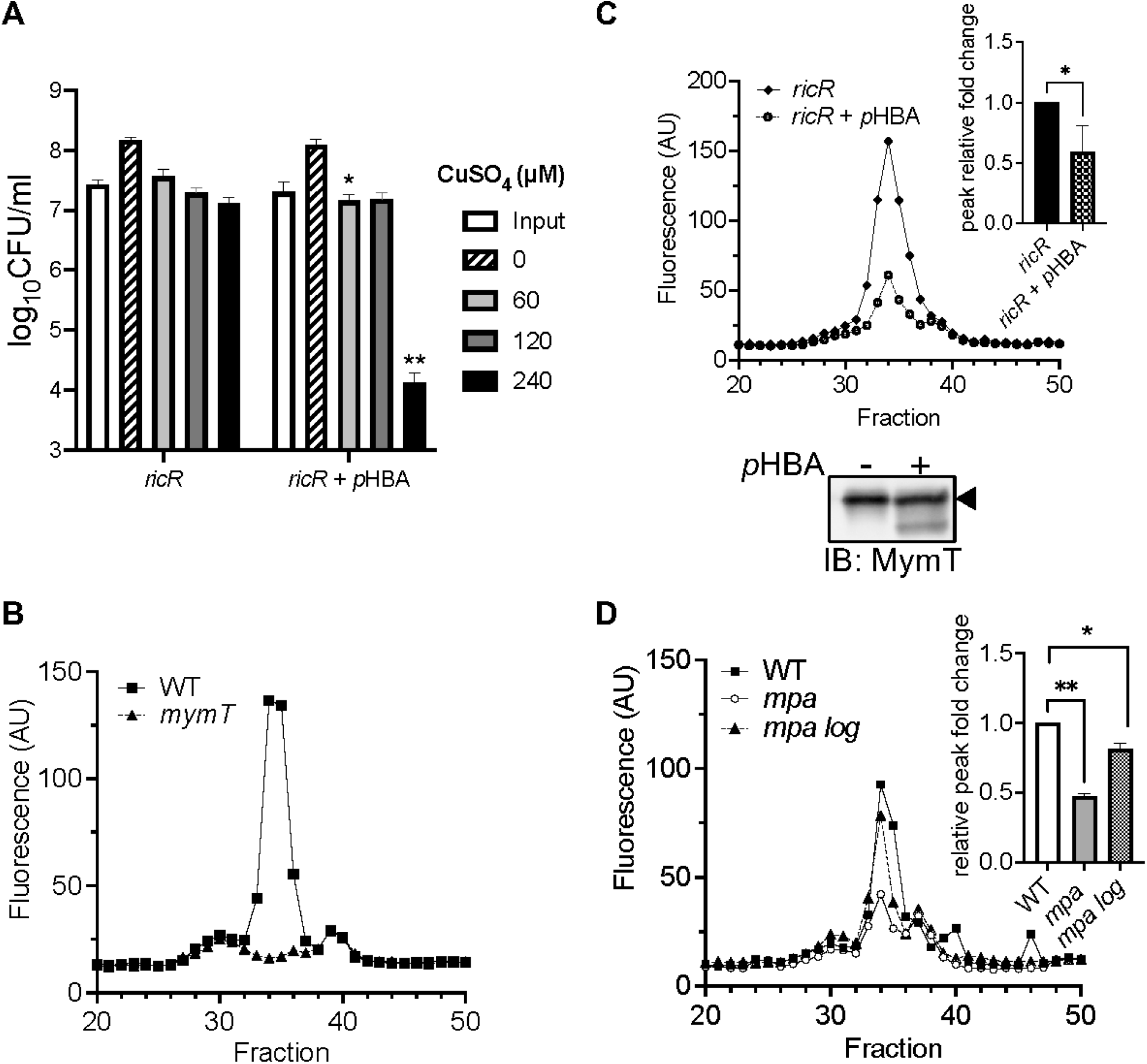
MymT activity is reduced in the presence of *p*HBA. (**A**) Cu sensitivity assay of a *ricR* mutant after preincubation with *p*HBA. Data are representative of two experiments, each done in technical triplicate. (**B**) MymT Cu(I)-thiolate luminescence (excitation = 280 nm, emission = 595 nm, cutoff = 325 nm). WT *M. tuberculosis* lysates exhibit a peak in luminescence in fraction 34, which is abolished in lysates of a *mymT* mutant. (**C**) Fractionated lysates from *M. tuberculosis ricR* mutant strain treated with or without *p*HBA. Inset: quantification of fold change. Lower section: Immunoblot for MymT of pooled fractions corresponding to the maximal fluorescence peak. Arrowhead indicates full-length MymT. (**D**) Fractionated *M. tuberculosis* lysates from WT, *mpa*, and *mpa log* strains. Inset: quantification of fold change. All quantifications were analyzed for significance using an unpaired t-test with * =*p* < 0.05; ** = p < 0.01. Unlabeled bars did not show significant differences.

Given that the metallothionein MymT plays a dominant role in Cu resistance in *M. tuberculosis*, we hypothesized *p*HBA could affect the function of MymT. Metallothioneins are found in all domains of life and protect cells from the potentially harmful effects of free Cu (22). MymT uses cysteines to bind reduced Cu [Cu(I)] (19). Given that aldehydes can readily form hemi-acetals with cysteines and disable protein function (23), we hypothesized *p*HBA reacts with cysteines in MymT, preventing its ability to bind and sequester toxic Cu. Cu-metallothioneins form solvent-shielded Cu(I)-thiolate cores that luminesce when excited by ultra-violet (UV) light (19, 24). Thus, a reduction in luminescence would indicate decreased Cu binding.

We first measured MymT Cu-thiolate cores from WT *M. tuberculosis* by fractionating whole-cell lysates of Cu-treated bacteria and measuring luminescence emitted from each fraction after excitation by UV light. A sharp peak of luminescence was observed in the fractionation profile that could be attributed to MymT, given that this peak was absent from lysates of a *mymT* null mutant (**Table 1, Fig. 4B**). We next examined the effect of *p*HBA on MymT luminescence in the *ricR* mutant. We observed a reduction of the MymT luminescence peak in lysates collected from the *ricR* mutant treated with *p*HBA compared to that of untreated bacteria, strongly suggesting *p*HBA affected the ability of MymT to bind Cu (**Fig. 4C**). To ensure equivalent MymT levels in *ricR* mutant lysates treated with or without *p*HBA, we pooled and concentrated the three fractions corresponding to the maximal MymT luminescence peak for immunoblot analysis. We found MymT levels were high and equivalent in *ricR* mutant fractions, whether or not they were *p*HBA-treated (**Fig. 4C**, lower panel). Interestingly, we consistently detected a smaller species of MymT in the *p*HBA-treated samples. It is possible this smaller species was caused by a direct modification of MymT by *p*HBA, an idea that remains to be tested.

We also tested if MymT luminescence was reduced in an *M. tuberculosis* PPS mutant compared to in the parental strain. The MymT luminescence peak in the *mpa* strain was significantly lower, and this reduction was restored to near WT levels in the *mpa log* double mutant (**Fig. 4D**). These data support the hypothesis that the accumulation of *p*HBA reduced either the amount or activity, or both, of MymT in a PPS mutant.

### A metabolic aldehyde sensitized *M. tuberculosis* to Cu

A recent hypothesis put forward by the Darwin and Stanley labs proposes host cell-derived aldehydes of metabolism contribute to bacterial control during infections (25). Macrophages undergo an increase in aerobic glycolysis, known as the Warburg Effect, following infection with *M. tuberculosis* and other pathogens (26–29). A by-product of aerobic glycolysis is MG, also known as pyruvaldehyde, which has been detected at millimolar concentrations in *M. tuberculosis-infected* mouse macrophages (30). Unlike with *p*HBA, MG pretreatment resulted in increased Cu resistance, leading us to hypothesize that MG preincubation specifically induced either an aldehyde or Cu resistance pathway, an idea we are testing. Nonetheless, when simultaneously added to bacteria with Cu, MG at normally non-toxic levels synergized with Cu to robustly kill *M. tuberculosis* (**Fig. 5A**, right panel, gray and black bars). Furthermore, MG treatment reduced luminescence of MymT in a *ricR* mutant (**Fig. 5B**). Overall, this result indicates that a physiologic aldehyde that is present during infections has the potential to sensitize *M. tuberculosis* to Cu by disrupting one or more Cu-responsive proteins.

**Fig. 5.**
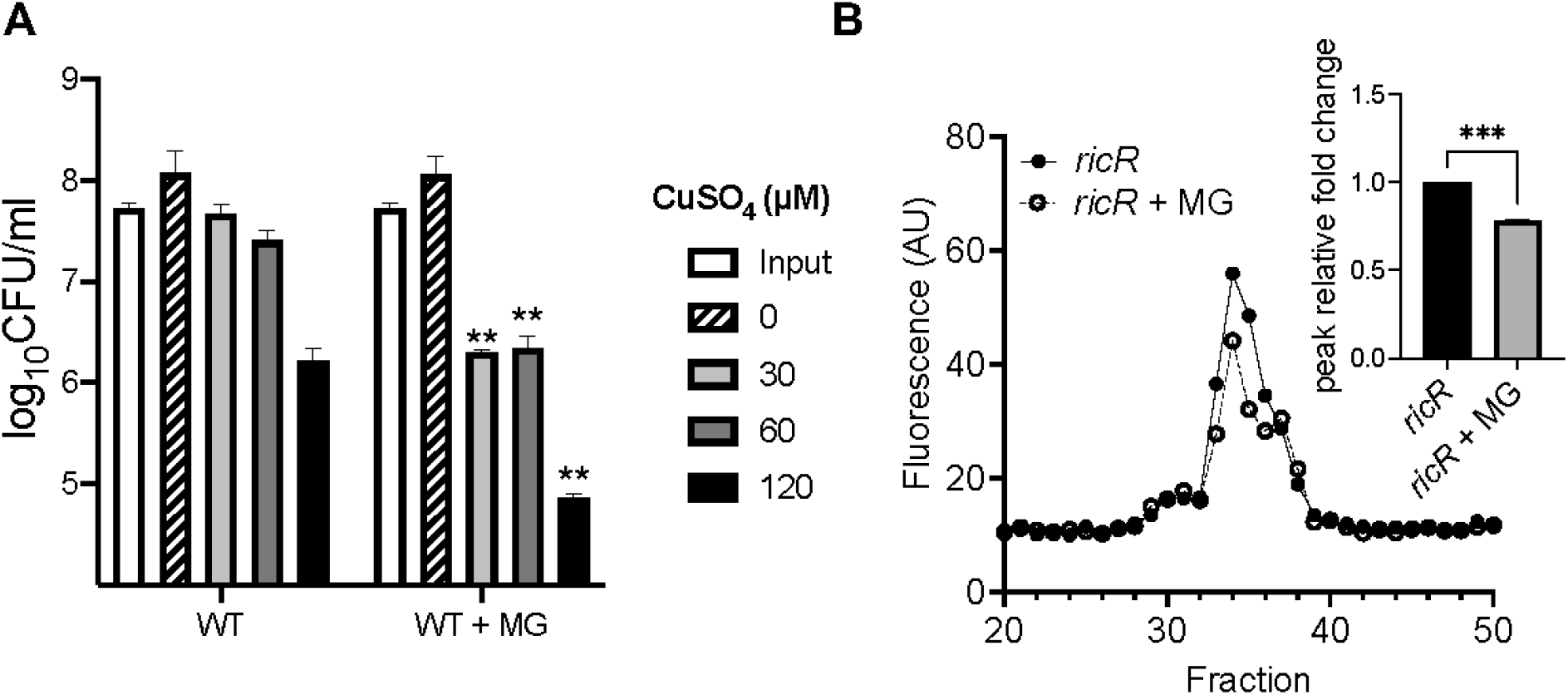
Methylglyoxal sensitizes *M. tuberculosis* to copper and disrupts MymT Cu-binding. (**A**) Cu sensitivity assay with the addition of MG to WT bacteria at day 0. Significant difference calculated comparing with the first strain on the x-axis at the same CuSO_4_ concentration using an unpaired t-test with *** =*p* < 0.001. (**B**) MymT Cu(I)-thiolate luminescence with a *ricR* mutant *M. tuberculosis* treated with or without MG. Inset: quantification of peak fold change relative to untreated, analyzed for significance using a two-tailed t-test (*p* < 0.05). Data are representative of two experiments, each done in technical triplicate.

## DISCUSSION

In this work, we sought to test if the Cu-sensitive phenotype of an *M. tuberculosis* PPS mutant was due to an accumulation of the aldehyde *p*HBA. We found that elimination of the PPS substrate Log, which is the source of *p*HBA, restored Cu resistance to WT levels. Furthermore, addition of *p*HBA to WT *M. tuberculosis* cultures was sufficient to sensitize bacteria to Cu. Cu sensitization in both PPS mutants and *p*HBA-treated WT *M. tuberculosis* was likely due to the reduced expression of genes needed for Cu resistance. We also showed that the Cu-binding function of MymT was altered in the presence of *p*HBA, possibly by disrupting cysteines in the protein. Finally, we showed that MG, an aldehyde produced by activated macrophages, also sensitized *M. tuberculosis* to Cu *in vitro*. Collectively, we propose a model whereby *p*HBA and other aldehydes can directly or indirectly disable Cu-sensing by RicR, leading to the constitutive repression of the RicR regulon, and also disrupt Cu-binding by MymT, preventing its ability to confer Cu resistance (**Fig. 6**).

**Fig. 6.**
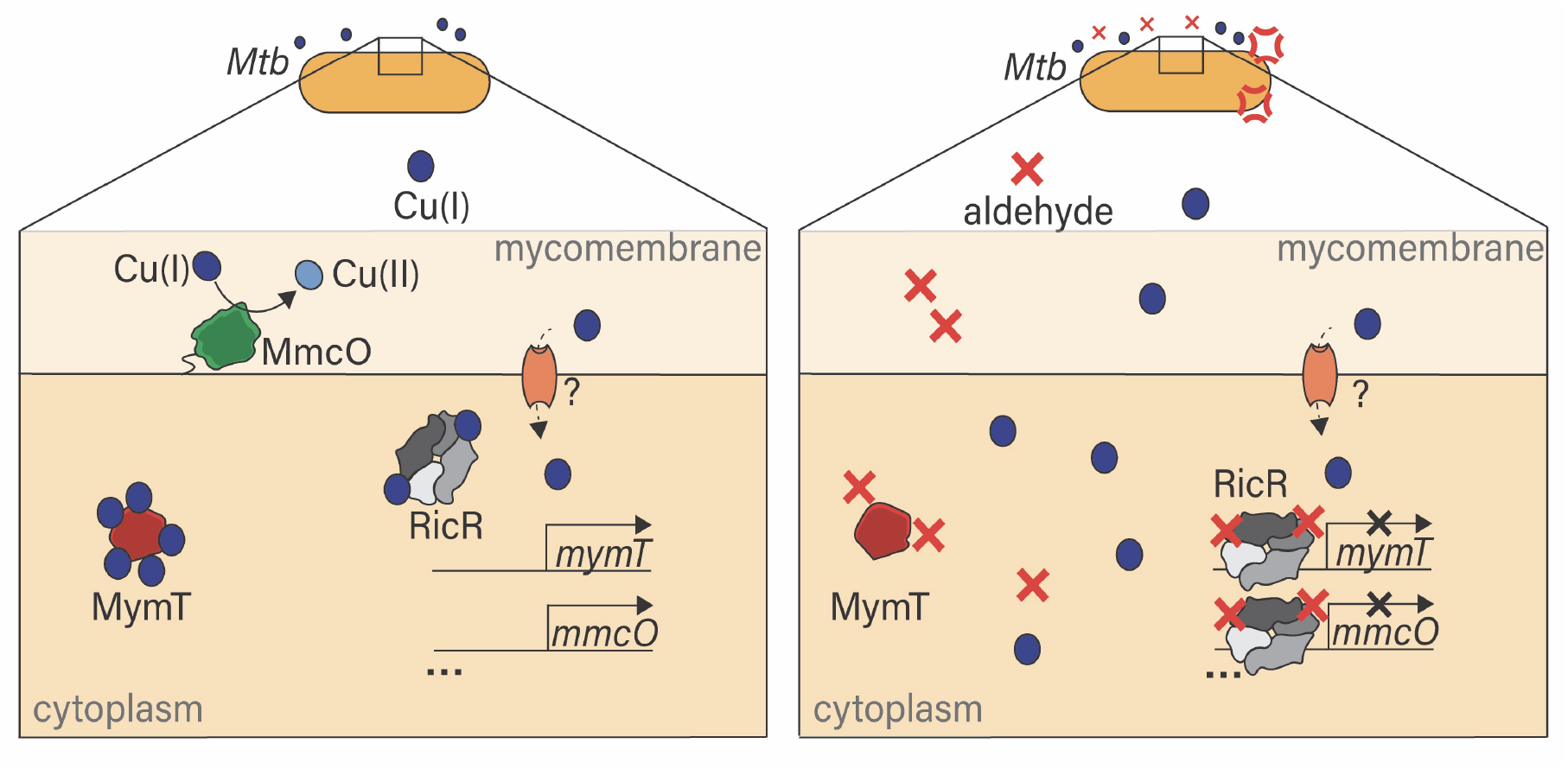
Proposed model of aldehyde sensitization of *M. tuberculosis* to copper. Left: in the absence of aldehyde, Cu(I) enters the cytoplasmic space through unknown transporters. Cu prevents RicR from binding to DNA, allowing expression of the RicR regulon genes, some of which mitigate Cu(I) toxicity, MymT sequesters Cu(I) to prevent it from damaging the cell, and MmcO is a periplasmic multicopper oxidase that oxidizes Cu(I) to its less toxic Cu(II). Right: in the presence of aldehyde, RicR cannot release from DNA, even in the presence of Cu, leading to repression of the RicR regulon and the observed Cu sensitivity of aldehyde-treated and PPS mutant *M. tuberculosis* strains. Additionally, MymT is unable to bind Cu in the presence of aldehyde.

In this study we also showed that a transposon disruption in *glpK*, which encodes a glycerol kinase, suppressed the Cu sensitive phenotype of a PPS mutant. This mutation also suppresses NO sensitivity (5). Along these lines, several independent studies recently reported that *glpK* mutations are often found in clinical *M. tuberculosis* isolates (31–33). These mutations are reversible and confer decreased susceptibility to antituberculosis drugs, suggesting phase variation occurs in response to antibiotic pressure on the bacteria. The Alland group proposed that the activation of a general stress response following a block in glycerol metabolism provides increased antibiotic tolerance (31). Similar to our hypothesis that a *glpK* mutation reduces aldehyde levels in PPS mutants, the Sassetti lab proposed that a block in glycerol metabolism, which produces MG, could protect bacteria by lowering endogenous aldehyde burden; this idea suggests aldehydes could sensitize bacteria to antibiotics (32).

In addition to the RicR regulon, the metal-responsive CsoR operon and Zur regulon were similarly regulated between *p*HBA-treated WT bacteria and PPS mutants. CsoR represses the expression of *ctpV*, which is a cation transporter involved in Cu resistance (17) and is repressed in PPS mutants and WT bacteria incubated with *p*HBA. CsoR and RicR are paralogs that use cysteines to sense Cu (13, 18); thus, it is possible that *p*HBA also disrupts the ability of CsoR to bind Cu, resulting in the constitutive repressed production of CtpV, a cation transporter implicated in Cu resistance (17).

The Zur regulon was upregulated in *p*HBA-treated *M. tuberculosis* (**Table 3**) and in PPS mutants (15). Unlike the Cu regulators, Zur is a zinc-responsive regulator that binds to and represses its promoters in the presence of excess zinc (34). Zur regulated genes are thus induced in low zinc and implicated in zinc uptake (34). The ESAT-6 cluster 3 (ESX-3) genes in the Zur regulon are also regulated by IdeR (iron dependent repressor) (35). However, no other IdeR-dependent genes were differentially expressed between untreated and *p*HBA-treated *M*. *tuberculosis*, similar to what we observed in PPS mutants (15). Notably, unlike the Cu regulators and Zur, IdeR does not use cysteine to coordinate iron.

In addition to the Zur regulon, several genes were induced in *p*HBA-treated *M. tuberculosis* and not in PPS mutants (**Table 3**). Some of these genes, e.g., *mesT* and Rv0195, are upregulated in hypoxia or after damage to the mycomembrane (36–38). It is possible these genes are specifically induced in response to extracellular aldehyde exposure and not by endogenously-produced aldehyde. Alternatively, the amount of *p*HBA we used for RNA-Seq cultures induced a transcriptional response that would not be achieved by what are likely much lower *p*HBA concentrations found in PPS mutants.

We also observed several uncharacterized operons with gene expression patterns shared between PPS mutants and *p*HBA-treated WT bacteria, including: the *cysDNC operon*, which encodes a sulfate-activating enzyme complex that is implicated in virulence, oxidative stress, sulfate limitation, and sensing of exogenous cysteine (39); Rv0762c-Rv0771, which includes a gene for a putative aldehyde dehydrogenase (*aldA*) and is predicted to be regulated by a probable ArsR-like metalloregulatory transcriptional repressor (Rv0576) (40, 41); and, Rv3249c-Rv3252c that is predicted to be controlled by WhiB4, a redox-responsive, iron-sulfur cluster-containing transcriptional repressor (**Table 2**) (42). Rv3249c-Rv3252c encodes the rubredoxins RubA and RubB and a putative monooxygenase AlkB. Relevantly, many of these genes are predicted to be regulated by metal-binding proteins that use cysteines to coordinate their respective metals (41, 42). Because aldehydes can form adducts with thiols in proteins, it is possible that cysteine-dependent metal coordinating proteins are particularly sensitive to aldehyde exposure.

Our data also suggest that aldehydes have a direct effect on the function of MymT. We observed that *p*HBA and MG reduced MymT luminescence, suggesting these aldehydes directly disrupted Cu-binding to MymT. An alternative explanation is that *p*HBA quenched MymT luminescence rather than directly preventing MymT Cu-binding; however, a smaller species of MymT formed only after *p*HBA treatment, suggesting this aldehyde directly affected the structure of MymT. This smaller species of MymT may have formed a more compact conformation due to a covalent interaction with *p*HBA, allowing it to migrate through SDS-PAGE gels more quickly. Alternatively, *p*HBA-modified MymT could have adopted a conformation that exposed it to peptidases, resulting in partially-degraded MymT. Unlike MymT, we could not test whether or not *p*HBA directly affected MmcO activity given that MmcO is a membrane-anchored protein without an established activity assay. Although MmcO is not predicted to use cysteines to function, its activity may nonetheless be affected by aldehydes, a hypothesis that remains to be tested.

A recent study by the Glickman lab identified an integrated system involving Rip1 protease and the PdtaS/R two-component system in *M. tuberculosis* that senses and mediates resistance to Cu and NO (43). The NO sensitivity of a *rip1* mutant is attributed to a block in chalkophore biosynthesis. Chalkophores bind to Cu with high affinity (43–45), and a follow up study supports a model that mycobacteria require chalkophores to acquire Cu during low Cu-conditions, i.e., this system is not for Cu resistance *per se* but for Cu acquisition (46). While it remains to be determined how Cu and NO resistance is conferred by Rip1, it is unlikely that aldehydes are involved given that known Cu-resistance genes are not repressed in a *rip1* mutant. Importantly, these data suggest there are additional ways for *M. tuberculosis* to be sensitized to NO and Cu.

An active anti-microbial role for aldehydes *in vivo* has yet to be established. In macrophages, aldehydes are produced at low levels during cellular metabolism (47, 48), but a shift to aerobic glycolysis following infection with *M. tuberculosis* leads to an increase of aldehydes such as glyceraldehyde-3-phosphate and MG (26–28, 30, 49–51). Induction of aerobic glycolysis plays a role in infection control given that the inhibition of this pathway in mouse macrophages leads to loss of some IFN-γ-dependent control of *M. tuberculosis* growth (26). Thus, aldehydes produced during this shift to glycolysis might contribute to antibacterial activity for the host.

More broadly, aldehydes may also have a role in the defense against other microbes. For example, a recent report by the Portnoy lab showed that in *Listeria monocytogenes (L. monocytogenes*), MG activates transcription of glutathione (GSH) synthase-encoding gene *gshF*, leading to increased GSH production and thereby activating the master virulence regulator PrfA (52). A *L. monocytogenes* mutant that lacks a gene encoding glyoxalase A, a key enzyme in methylglyoxal detoxification, has decreased GSH levels *in vitro* and is attenuated for infection. Together these results suggest that MG is an important cue for some pathogens to turn on virulence programming.

Our data provide the first evidence that aldehydes can sensitize *M. tuberculosis* to Cu. Importantly, despite well-established evidence of their toxicity, the antibacterial mechanisms of aldehydes are relatively uncharacterized; thus, our study may begin to provide new insight into how aldehydes can target and inactivate bacterial pathways needed for survival.

## MATERIALS AND METHODS

### Bacterial strains, growth conditions, plasmids, and primers

The bacterial strains, plasmids, and primers used in this work are listed in Table 1. *M. tuberculosis* strains were grown in 7H9 liquid media (Difco) supplemented with 0.2% glycerol, 0.05% Tween 80, 0.5% bovine serum albumin (BSA), 0.2% dextrose, and 0.085% sodium chloride (ADN) (referred to as “7H9c” from here on) or Sauton minimal media (3.7 mM potassium phosphate, monobasic; 2.4 mM magnesium sulfate; 30 mM L-asparagine; 3.5 mM zinc sulfate; 9.5 mM citric acid; 6.0% glycerol; 0.005% ferric ammonium citrate; 0.05% Tween-80). Cultures were grown at 37°C without agitation in vented flasks (Corning). For *M. tuberculosis* growth on solid media, Middlebrook 7H11 agar (Difco and Remel) was supplemented with Middlebrook OADC (oleic acid, albumin, dextrose, and catalase; BBL). *M. tuberculosis* strains were grown in 50 μg/ml kanamycin, 50 μg/ml hygromycin when necessary.

For CuSO_4_ solutions: stock solutions were made by dissolving the appropriate amount of CuSO_4_ powder (Fisher Scientific) in water and filter-sterilizing with a 0.45 μ filter. For aldehyde solutions, 50 mM *p*HBA was made with *p*HBA powder dissolved in water and filter-sterilized using a 0.45 μ filter. MG (5.5 M) was diluted in sterile water just before use. Both aldehydes were purchased from Sigma-Aldrich, Inc.

### RNA-Seq

*M. tuberculosis* cultures were grown in 7H9c media and treated with 1.2 mM *p*HBA at an optical density at 580 nm (OD_580_) of 1.0 (late logarithmic to early stationary phase) or left untreated. 24 hours later, RNA was purified as previously described (15). RNA was isolated from three biological replicate cultures. Library preparation and Illumina HiSeq Sequencing were performed by GENEWIZ, LLC. Sequence reads were mapped to the *M. tuberculosis* H37Rv genome sequence (RefSeq identifier GCF_000195955.2) using bwa v0.7.17 (53) and sorted using samtools v1.9 (54). An average of 97.8% of reads were mapped to the reference genome, indicating high quality of samples. Given a locus of interest, *socAB*, was not included in the assembly annotation from RefSeq, we added it to the annotation files at genomic coordinates NC_000962.3:1,933,937-1,934,497. Using the alignment files generated for each sample, the *featureCounts* command in Subread v2.0.1 (55) was used to count the reads mapping to each gene in the reference. Read counts per gene and sample (feature counts) were loaded into R v4.2.0 (56) for further analysis using the package DESeq2 v1.36.0 (57). DESeq2’s normalization function was used to normalize the counts to make expression levels more comparable between the different samples. To compare gene expression changes between *p*HBA-treated and untreated samples, the Wald test was used to generate *p*-values and log_2_-fold changes. Genes with an adjusted *p*-value of <0.05 and fold change of >2.0 when comparing *p*HBA-treated to untreated *M. tuberculosis* were considered differentially-expressed genes (**Fig. 2** and **Table S1**). Raw sequencing data files are available in a PATRIC public workspace: (https://www.bv-brc.org/workspace/ginalimon@bvbrc/Limon_Darwin_RNA-Seq). The volcano plot was generated using python v3.10.6 and package bioinfokit v2.1.10 (58),

### Cu-sensitivity assay

Cu-sensitivity assays were performed as previously described (15, 18). Briefly, *M. tuberculosis* strains were grown in 7H9c to an OD_580_ = 0.5-1.0. Bacteria were washed once with Sauton minimal media with no added Cu and collected using a low-speed centrifugation (150 *g*) to remove clumped cells. Supernatants containing mostly unclumped bacteria were diluted to OD_580_ = 0.08 in Sauton minimal media. 194 μl of this diluted culture were transferred to wells of 96-well plates; 6 μl of the appropriate stock concentration of CuSO4 or aldehyde or both were added to the desired final concentration. Plates were incubated at 37°C for 10 days, after which cultures were diluted and inoculated onto 7H11 agar Y-plates. Plates were incubated for 14-21 days before enumerating CFU. As previously reported (15), we used a range of CuSO4 concentrations due to variability of Cu-sensitivity between experiments. Each experiment was done at least twice in technical triplicate.

### *M. tuberculosis* lysate preparation for immunoblotting

For all blots, bacteria were grown in Sauton minimal media with no added Cu. For MmcO blots, at OD_580_ = 0.5-1.0, cultures were treated with 1.2 mM *p*HBA for 4 hours before treatment with 50 μM CuSO4. Bacteria were harvested 24 hours after the addition of CuSO4. Bacterial densities were measured and equivalent cell numbers were collected based on the OD_580_ of the cultures. For example, an “OD_580_ equivalent of 1” indicates the OD_580_ of a 1 ml culture is 1.0. For most assays, 5 OD_580_ units were collected and washed once with phosphate-buffered saline (DPBS, Corning, Inc with 0.05% Tween-80) to remove BSA in 7H9c media. Bacterial pellets were then resuspended in 300 μl TE buffer (100 mM Tris-Cl, 1 mM EDTA pH 8.0), and transferred to bead-beating tubes with 200 μl zirconia beads; tubes were beaten for 30 seconds three times, with icing for 30 seconds in between in a mini bead beater (all materials from Bio-Spec). 150 μl of lysate was transferred into new tubes with 50 μl 4× SDS sample buffer (250 mM Tris pH 6.8, 2% SDS, 20%β-mercaptoethanol, 40% glycerol, 1% bromophenol blue) and boiled at 100°C for 10 minutes. Proteins were separated by sodium dodecyl sulfate polyacrylamide gel electrophoresis (SDS-PAGE) and transferred onto nitrocellulose membranes. Membranes were blocked in 3% milk or BSA prior to incubation with polyclonal rabbit antibodies as indicated in the figure legends. For loading controls, the same membranes were stripped with 0.2 N NaOH as described elsewhere (59) and re-blocked before incubation with antibodies to *M. smegmatis* PrcB (60).

### Construction of a Δ*ricR::hyg* mutant

We made an *M. tuberculosis ricR* deletion mutant strain using a previously described method (15). Briefly, pYUB854 (61) was used to clone sequences encompassing ~700 bp upstream (5’) and ~700 downstream (3’) of the *ricR* gene. The 5’ and 3’ sequences, including the start and stop codons, respectively, were cloned to flank the hygromycin resistance cassette in pYUB854. The plasmid was digested with PacI, and approximately 1 μg of linearized, gel-purified DNA was used for electroporation into *M. tuberculosis. M. tuberculosis* strains were grown to an OD_580_ of ~0.4 to 1, washed, and resuspended in 10% glycerol to make electrocompetent cells as described in detail in (62). Bacteria were inoculated onto 7H11 agar with 50 μg/ml hygromycin as needed; a no-DNA control electroporation was done to control for spontaneously antibiotic-resistant mutants. Two weeks after plating, colonies were picked and inoculated into 200 μl 7H9c with antibiotics and then inoculated into 5 ml cultures for further analysis. Mutants were confirmed by PCR and sequence analysis.

### MymT luminescence from *M. tuberculosis* lysates

We adapted a previously reported protocol for measuring Cu(I)-thiolate core luminescence for use on filtered *M. tuberculosis* lysates (19). *M. tuberculosis* cultures were grown in Sauton minimal media to an OD_580_ = 0.3-0.5 and treated with *p*HBA to a final concentration of 1.2 mM as needed and 50 μM CuSO_4_ four hours later. 24 hours after Cu addition, 12 OD_580_ equivalent cell numbers were harvested by centrifugation and washed twice with buffer (10 mM HEPES, 150 mM NaCl pH 7.4). Bacteria were resuspended in 700 μl of the same buffer and lysed by bead beating as described for preparing lysates for immunoblotting. Lysates were then centrifuged for 7.5 min at 20,000 *g* and the supernatants were passed through a 0.2 μ spin filter twice before application onto a Superose-6 10/300 GL column (Cytiva). Fractions were transferred to a UV grade 96-well plate (Corning) and luminescence was measured with excitation at 280 nm, emission 595 nm, and a cutoff of 325 nm.

For immunoblotting proteins in fractionated lysates, 200 μl of fractions corresponding to the three at the peak fluorescence collected using method above were frozen at −20°C before being pooled and concentrated in a 0.5 ml centrifugal filter (Amicon). Samples were boiled in SDS sample buffer for 10 minutes before separation on 15% SDS-PAGE gels and transferred onto nitrocellulose membranes. Polyclonal MymT antibodies used for immunoblotting were a kind gift from Ben Gold and Carl Nathan (19).

## ACKNOWLEDGEMENTS

We thank S. Becker, S.A. Stanley, P. Tran and J.H. Yoo for reviewing a draft version of this manuscript. We thank X. Shi for making MHD799. This work would not have been possible without preliminary experiments by M. Samanovic-Golden, the technical expertise of J. Ilmain (V. Torres Lab), or advice from S. Kahne. We thank J. Belasco, K. Cadwell, and M. Pacold for helpful suggestions. This work was supported by NIH grant AI153197 awarded to K.H.D and S.A.S. We thank the Office of Science and Research High-Containment Laboratories at NYU Grossman School of Medicine for their support in the completion of this research.

---

**Supplemental Table S1. Gene expression data of transcripts of *M. tuberculosis* treated with and without *p*HBA**.

